# A green fluorescent protein for live imaging in hyperthermophiles

**DOI:** 10.64898/2026.03.09.710647

**Authors:** Yin-Wei Kuo, Arthur Radoux-Mergault, Tom Dubois, Alice Cezanne, Fan Zhang, Pier Andrée Penttilä, Michaela Wagner, Gautam Dey, Sonja-Verena Albers, Buzz Baum

## Abstract

Hyperthermophiles, organisms that thrive at temperatures above 60 °C, have played important roles in biotechnology and promise to reveal new biology. However, how these cells live remains poorly understood in part due to the lack of bright, thermostable fluorescent proteins that can be used to study protein localisation and dynamics at high temperatures. To overcome this challenge, here we describe the development of “Matcha”, a green fluorescent protein that we have engineered from Thermal Green Protein by directed evolution in the thermophilic archaeon *Sulfolobus acidocaldarius*. The screen identified 7 mutations that when combined led to an ∼50-fold increase in the brightness of Matcha *in vivo* at physiological temperatures. Since this is sufficient for live cell imaging, we were then able to use Matcha-fusion proteins to study the division ring dynamics in *Sulfolobus*. Remarkably, this analysis reveals that, while ESCRT-III rings are disassembled as cells complete division, CdvA forms a stable polymeric ring that persists, and is asymmetrically inherited by one of the two daughter cells following cytokinesis. This study highlights the power of Matcha as a tool to shed light on our understanding of the cell biology of hyperthermophiles.

## Introduction

The discovery of fluorescent proteins has transformed cell biology research over the past eight decades. It has enabled the visualization and tracking of cellular structures in living cells with unprecedented sensitivity and precision. However, the use of fluorescent proteins has been largely limited to model organisms that live in mesophilic environments. This has limited our understanding of the cell biology of life at high temperatures which has important implications in understanding how organisms adapt to the extreme environments and perhaps the origin and evolution of life on earth.

While earlier protein engineering efforts have reported several fluorescent proteins (FPs) with enhanced thermal stability *in vitro* (*1–6*), these have not been sufficiently bright to study the physiology of hyperthermophiles (*7, 8*). Thus, there remains a pressing need to develop a fluorescent protein that provides sufficient signal-to-noise ratio for live imaging of hyperthermophilic organisms. To fill this gap, we employed a directed evolution strategy to generate a highly improved green fluorescent protein and applied it to live imaging of proteins involved in cell division of the thermophilic archaeon *Sulfolobus acidocaldarius*.

## Results and Discussion

### Engineering of Matcha in a hyperthermophilic archaeon

Thermal green protein (TGP) (*1*) was recently reported to be visible when conjugated to a model substrate β-galactosidase (LacS) and expressed in *Sulfolobus acidocaldarius* (*8*) - a model hyperthermophilic archaeon that grows at an optimal temperature of ∼75 °C. To test the utility of TGP as a fluorescent tag for live cell imaging, we fused TGP to the N-terminus of several different protein-of-interests with a C-terminal hemagglutinin (HA) tag, including LacS, cell division protein, and DNA binding proteins. This set of fusion proteins was then expressed under the control of an arabinose-inducible promoter. Flow cytometry analysis was performed to assess green fluorescence signal in live cells, and immunostaining of HA-tag in ethanol-fixed cells was used to examine the expression level of each construct. After a long (overnight) induction at 75 °C, the only green fluorescence signal that could be detected above the empty vector control was in the cells expressing Matcha fused to LacS (Fig. S1), which is a highly stable protein (*9*). Even in this case, the portion of GFP positive cells was much smaller (∼30%) than the HA positive cell population measured from the ethanol-fixed cells (∼94%), indicating a large proportion of the TGP-LacS molecules in each cell were not fluorescent under these conditions. This fluorescence was partially enhanced with reduced temperature (Fig. S2A) at which we confirmed that Sulfolobus can still grow and divide with a slower rate (Fig. S3). However, even at lower growth temperatures, little fluorescent signal could be seen after a 4-hour induction (about half of the cell cycle length at 65 °C; Fig. S2B). For all these reasons, in its current form TGP cannot be used as a tool for fluorescent detection of protein-of-interests in *Sulfolobus*.

To improve the *in vivo* fluorescent properties of TGP, we used a directed evolution approach. For the screen, we generated a library that encoded a complete set of single amino acid variants (i.e., a site-saturation variant library) at 47 selected sites in the TGP protein. The modest library size (∼10^3^ unique clones) compensated for the relatively low transformation efficiency of *Sulfolobus* (∼10^4^ clones per transformation) compared to the commonly used cell models like *E. coli*. We then fused the entire set of mutants to LacS, transformed them into cells and selected for the brightest clones by successive rounds of flow cytometry, sorting, and sequencing (Fig. 1A). In this way we identified 7 independent mutations (Fig. 1B, 1C, Table S1; full sequence in Fig. S4) that, when combined, led to a 47±7-fold (mean±SD unless otherwise noted) enhancement in the normalised fluorescent signal at 75 °C (Fig. 1D, 1E).

**Figure 1.**
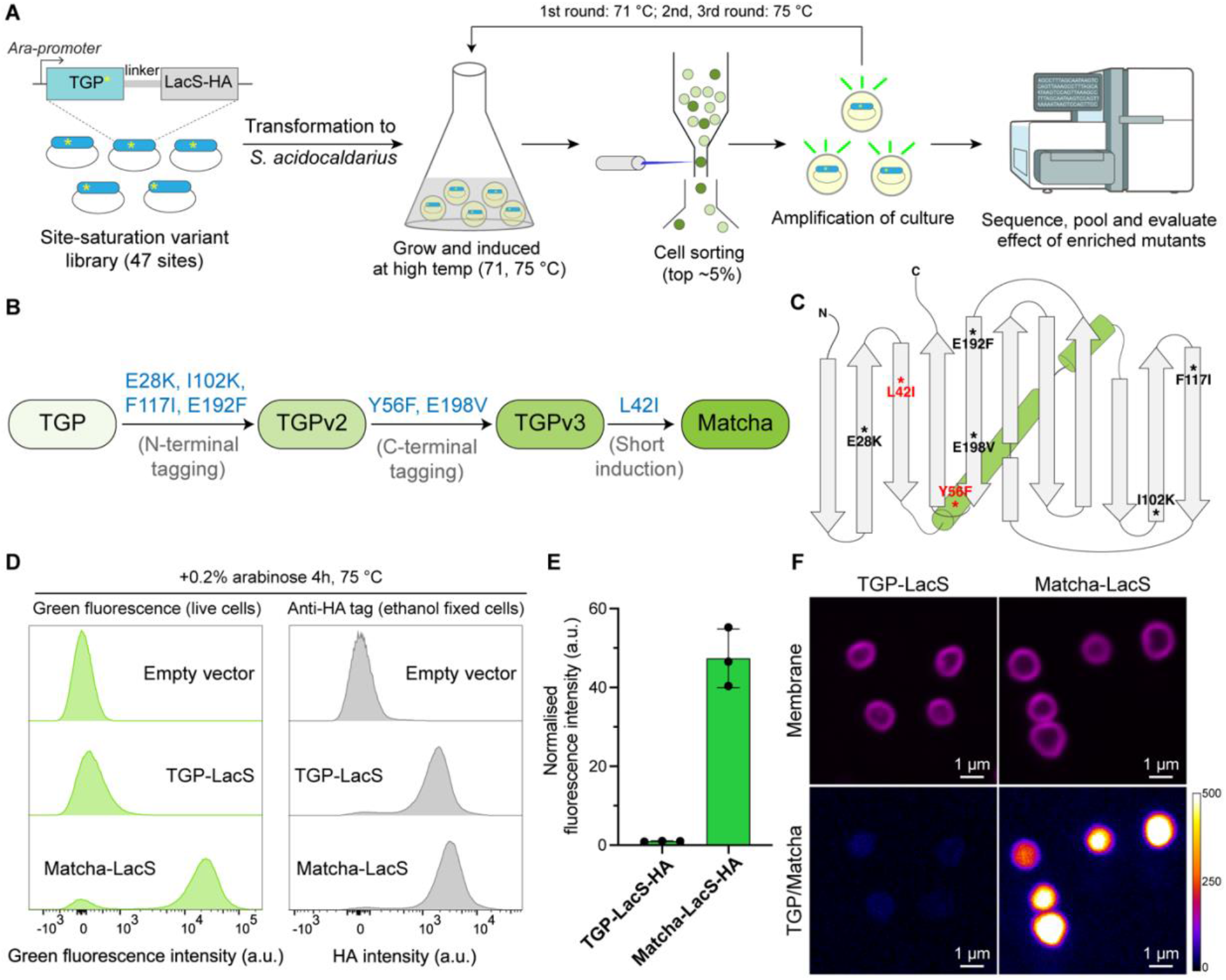
Development of Matcha as a fluorescent protein tag in *S. acidocaldarius*. **(A)** Schematic process of the directed evolution strategy to improve *in vivo* fluorescence using TGP as a starting template. **(B)** Engineering of Matcha by sequential incorporation of mutations to TGP. **(C)** The 7 mutation sites used for the evolution of Matcha. Residues located inside the β-barrel are marked in red. The secondary structure map is based on PDB 4TZA (*1*). **(D)** Example histograms of flow cytometry analysis of live cell green fluorescence signal (left) and HA (right) signal for expression level normalization (immunostained fixed cells). **(E)** Normalised green fluorescence signal of cells expressing LacS fused to TGP or Matcha respectively at 75 °C (4 hr arabinose induction) measured by flow cytometry. The normalized fluorescence signal was measured by dividing the average green fluorescence signal of live culture with the average anti-HA signal measured from the corresponding ethanol-fixed cells, and rescaled by the mean of TGP-LacS experiments (N=3 biological replicates each). **(F)** Example SoRa spinning disk confocal images of cells expressing LacS fused to TGP or Matcha at 75 °C (4 hr induction). Intensity of the green fluorescence channel (bottom) was shown by the colour scale bar.

We named this new green fluorescent protein ‘Matcha’ after the hot beverage with a characteristic green colour. Consistent with our flow cytometry analysis, cells expressing Matcha-LacS were significantly brighter than those expressing TGP-LacS at 75 °C when imaged using a spinning disk super-resolution microscope (SoRa) (Fig. 1F).

We followed up this analysis with a biophysical characterisation of the protein. Recombinant Matcha purified from *E. coli* was found to have similar absorption and emission spectra (Fig. S5A, B) and a ∼45% increase in brightness compared to TGP (Fig. S5C). *In vitro* stability assays also showed that Matcha and TGP were both highly stable with a melting temperature above 90 °C and were both much more resistant to denaturation by guanidine hydrochloride than EGFP (Fig. S6).

### Matcha as a fluorescent tag for high temperature live cell imaging

To determine if Matcha is sufficiently bright to use in live cell imaging, we first imaged *Sulfolobus* cells expressing the chromatin protein, Cren7 (*10, 11*), fused to Matcha. At 75 °C, Matcha-Cren7 was sufficiently bright to label the nucleoids of cells at different stages of the cell cycle (Fig. 2A, Supplementary video 1).

**Figure 2.**
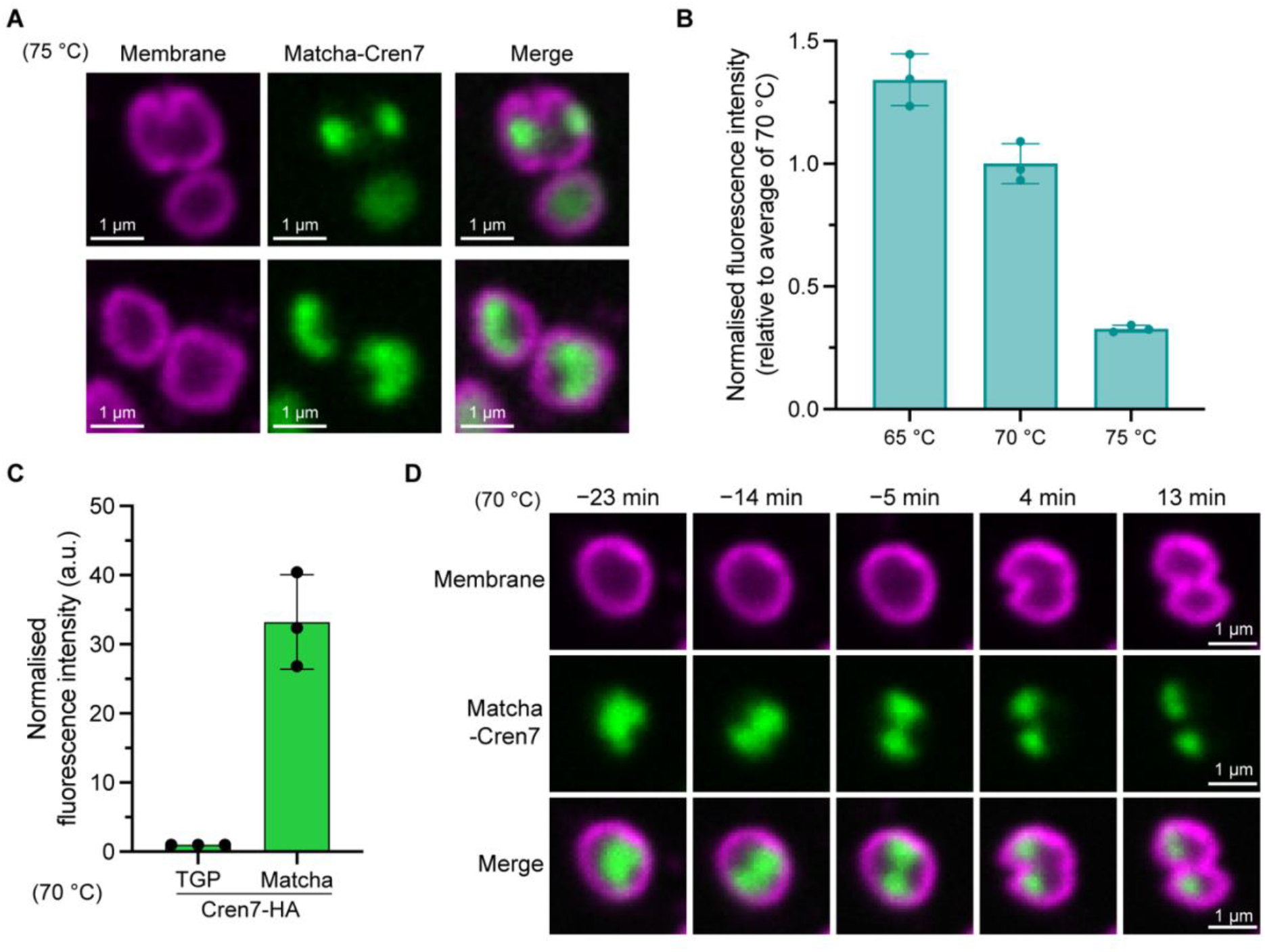
High temperature live imaging of chromatin binding protein Cren7 with Matcha. **(A)** Example images of cells expressing Matcha-Cren7 at 75 °C (4 hr arabinose induction). **(B)** Normalized fluorescence intensity of cells expressing Matcha-Cren7 from 65 to 75 °C (4 hr arabinose induction). The normalized intensities were rescaled relative to the average at 70 °C. N=3 biological replicates each. **(C)** Matcha-Cren7 showed stronger normalized fluorescence signal than TGP-Cren7 at 70 °C (after 4 hr arabinose induction). N=3 biological replicates each. **(D)** Example time-lapse images of a Matcha-Cren7 expressing cell during division. Time points are labelled relative to the onset of membrane constriction.

To examine if reduced growth temperature further enhanced the fluorescence, we measured the normalized fluorescence of cells expressing Matcha-Cren7 at different temperatures by flow cytometry. This revealed a 3.1±0.2-fold increase in fluorescence signal when decreasing the growth temperature from 75 to 70 °C. Lowering the growth temperature further to 65 °C, however, only led to a marginal 34±11% increase in normalized fluorescence intensity (Fig. 2B). Matcha-Cren7 also showed a strong fluorescence signal improvement compared to TGP-Cren7 at 70 °C (33±7-fold; Fig. 2C), consistent with our analysis of Matcha-LacS (Fig. 1E). Given the considerable enhancement in signal intensity and moderate increase in growth doubling time (∼60% longer than at 75 °C, Fig. S3A), we concluded that 70 °C would be ideal for timelapse fluorescent imaging of *Sulfolobus* using Matcha. Under these conditions, we were able to use Matcha-Cren7 to reveal nucleoid dynamics similar to those observed previously using a DNA binding dye (*12*). This includes a dynamic phase that precedes nucleoid compaction and segregation (Fig. 2D, Supplementary video 2). These results demonstrate the utility of Matcha for live cell imaging of *Sulfolobus*.

### Live cell imaging of ESCRT-III homologs during cell division using Matcha

Encouraged by the results obtained using Matcha-Cren7, we wanted to use Matcha to visualize the process of cytokinesis itself. Most current models of *Sulfolobus* cell division assume that CdvA protein forms a medial ring that recruits the ESCRT-III homolog CdvB. This CdvA/B ring then serves as a template for the assembly of a composite division ring composed of the two ESCRT-III homologs, CdvB1 and CdvB2. The sequential disassembly of CdvB, followed by CdvB1 and CdvB2 via the activity of Vps4 then drives membrane constriction and abscission (*13, 14*). Importantly, however, this sequence of events has been inferred from cells that have been fixed and stained using antibodies, never by live cell imaging.

To image the process live, we generated cells expressing CdvB, CdvB1 and CdvB2 fused to Matcha at their C-terminus respectively under the arabinose-inducible promoter. While CdvB-Matcha was bright enough to see rings in live cells (Fig. 3A), the fluorescence signal was dim and these rings were short lived - likely reflecting the low stability of CdvB as a rapidly degraded substrate of the proteasome (*9, 15*). As a result, we were unable to use CdvB-Matcha to image the dynamics of CdvB. However, we were able to use Matcha to follow the formation, constriction and disassembly of the medial CdvB1-Matcha and CdvB2-Matcha ring structures (Fig. 3B, 3C; Supplementary video 3, 4), which are more stable than CdvB (*9*).

**Figure 3.**
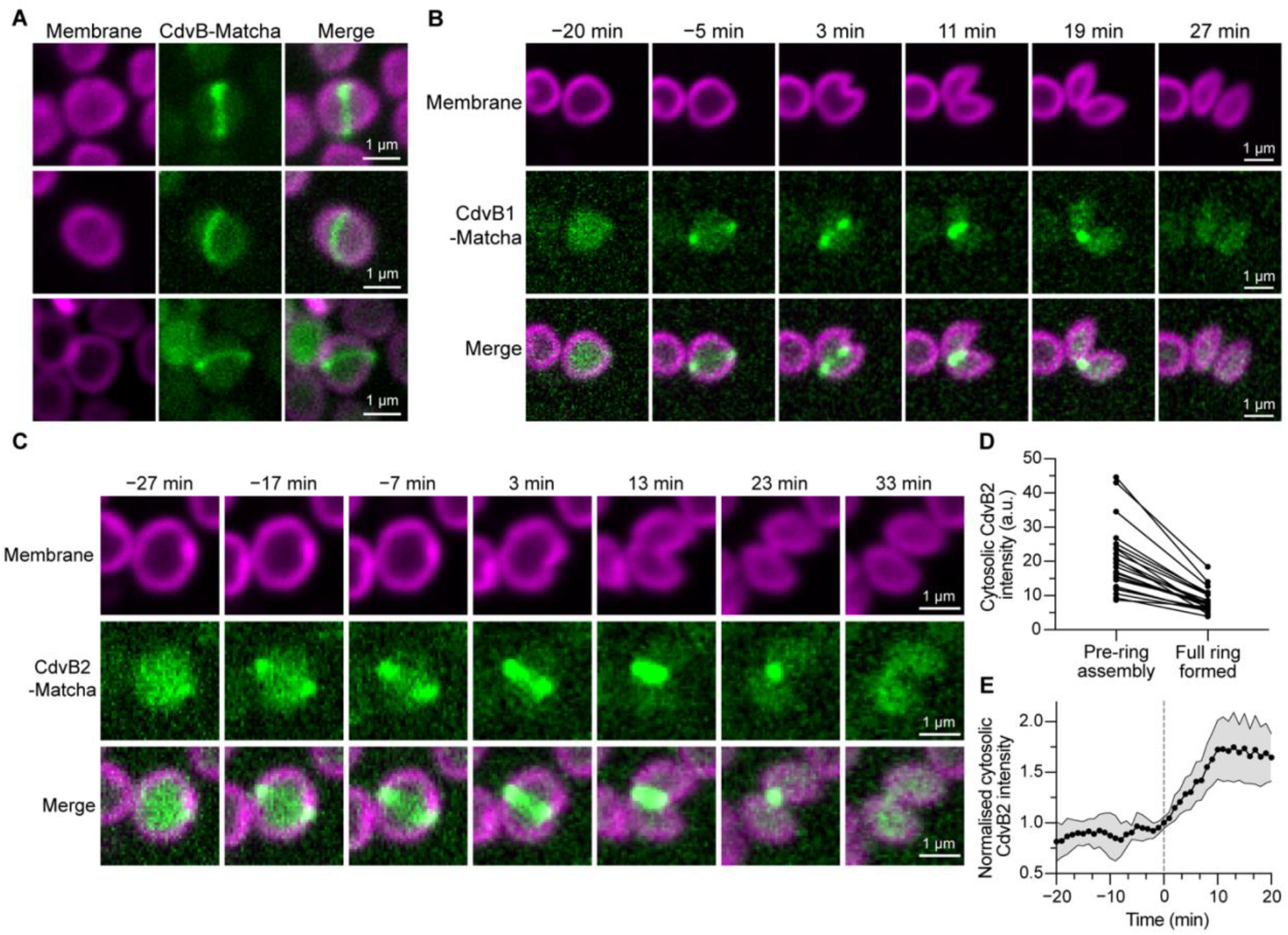
Visualization of archaeal ESCRT-III homologs with Matcha during cell division. **(A)** Example images of CdvB-Matcha ring at 70 °C (4 hr arabinose induction). **(B)(C)** Example timelapse images of dividing cells expressing CdvB1-Matcha (B) or CdvB2-Matcha (C) at 70 °C after ∼4 hr arabinose induction. Time points are labelled relative to the onset of membrane constriction. **(D)** Quantification of relative cytosolic CdvB2-Matcha signal before and after full ring assembly at 70 °C. N=23 cells from 3 biological replicates. **(E)** Time trace of normalized cytoplasmic CdvB2 signal at late stage of cytokinesis. The end of membrane constriction was set to the time=0 min point. The trace is averaged over 9 division events from 3 biological replicates (mean±SD).

Since CdvB2-Matcha gave the clearest signal, we used this for an analysis of the dynamics of ESCRT-III ring formation, contraction and disassembly. To better understand the polymerization and depolymerization dynamics, we first measured the cytosolic fluorescence of CdvB2-Matcha during cell division. This revealed that cytosolic signal of CdvB2-Matcha decreases as the ring forms, likely reflecting monomer polymerisation over a period of a few minutes (Fig. 3C, 3D). This decrease in cytosolic fluorescence accompanied by ring formation can also be observed in cells in which CdvB2-Matcha rings formed parallel to the imaging plane (Fig. S7A, S7B yellow bracket; Supplementary video 5).

After the CdvB2-Matcha ring had formed, constriction occurred progressively over a period of ∼30 minutes at 70°C. This constriction time was longer than previously observed (*16*), likely as a consequence of the lower temperature and the impact of Matcha-CdvB2 expression on the ring (as in seen in human cells expressing fluorescently tagged ESCRT-III proteins (*17–20*)). After membrane constriction was complete, we observed an increase in the cytosolic pool of CdvB2-Matcha (Figure 3E, Fig. S7C) prior to abscission. In most cases, the CdvB2 polymer disassembly began once rings had constricted to a point (Figure 3C, E). However, in the few rare cases in which cells failed division (perhaps due to CdvB2-Matcha expression), we observed CdvB2 structures that shortened over time as if depolymerizing from their ends instead of fragmenting prior to disassembly (Fig. S8).

### Live cell imaging of CdvA during cell division using Matcha

In contrast to the ESCRT-III homologs (CdvB, CdvB1, CdvB2), little is known about the role and fate of the archaeal-specific CdvA protein during cell division (*13, 14, 21*). To visualise CdvA dynamics during division, we next performed live imaging of cells expressing Matcha-CdvA under an arabinose-inducible promoter. The experiment was carried out at 65°C to reduce toxicity that appears to result from overexpression at high temperature. Unexpectedly, under these conditions Matcha-CdvA formed a ring that remained throughout the process of cytokinesis (Fig. 4A and 4B). As the cell constricted, so did the Matcha-CdvA ring. Strikingly, instead of being disassembled and degraded like its partner protein CdvB, Matcha-CdvA persisted through cytokinesis and was then asymmetrically inherited by one of the two daughter cells after division (Fig. 4A, 4C, Supplementary video 6).

**Figure 4.**
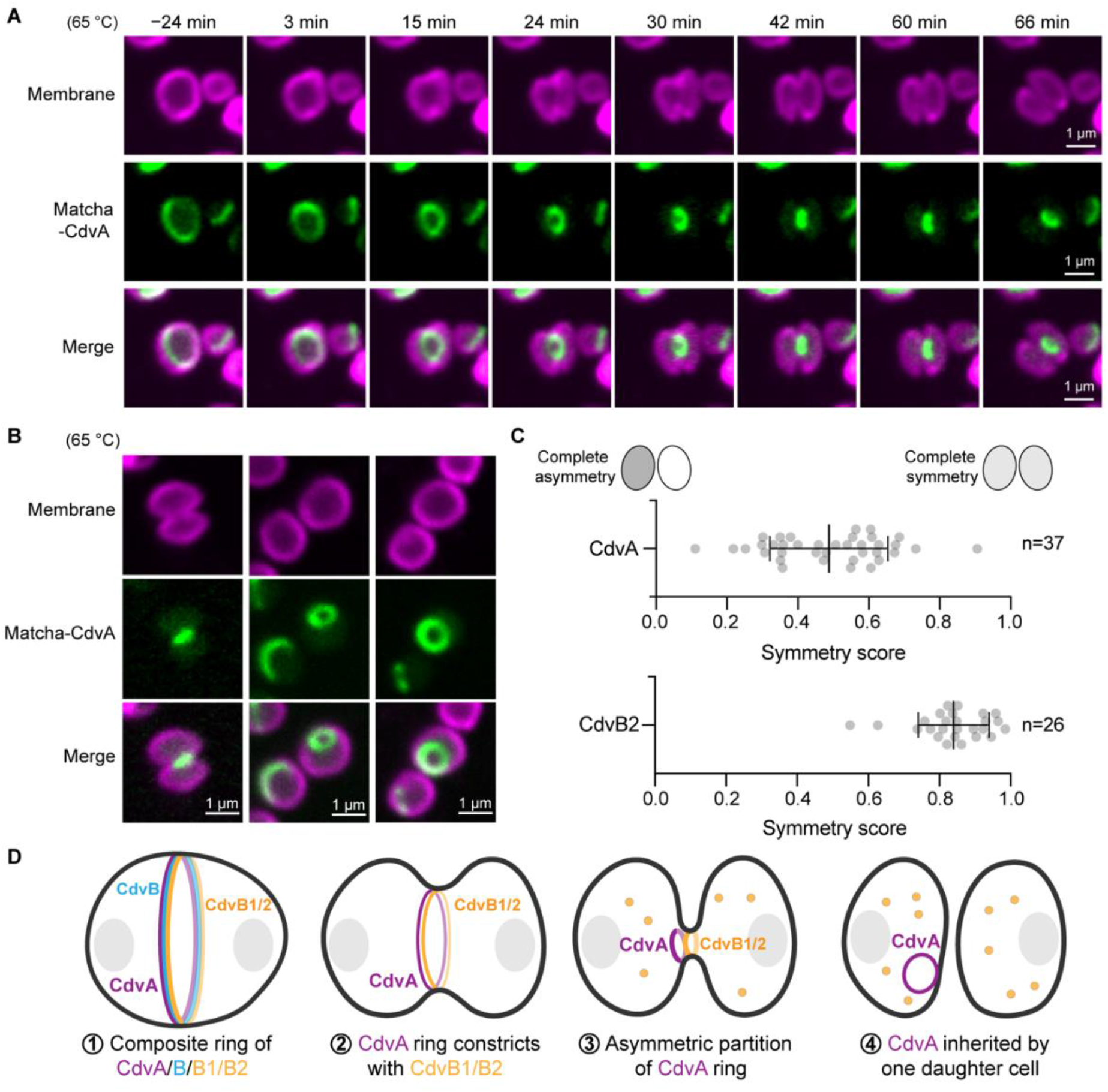
CdvA fused to Matcha forms ring polymer that persists during cytokinesis. **(A)** Example time-lapse images of cells expressing Matcha-CdvA by an arabinose-inducible promoter. Time points are labelled relative to the onset of membrane constriction. **(B)** Example images of Matcha-CdvA rings. **(C)** Quantification of CdvA and CdvB2 inheritance after cytokinesis, showing CdvA is inherited asymmetrically (symmetry score=0.49±0.17; mean±SD, n=37 events from 3 biological replicates) while CdvB2 inheritance is largely symmetric (symmetry score=0.84±0.10; mean±SD, n=26 events from 3 biological replicates). We defined the symmetry score as the ratio of the average CdvA or CdvB2 intensity in the two daughter cells (the lower intensity one divided by the higher one), where the score of 0 represents completely asymmetric and 1 represents completely symmetric inheritance. Note that CdvB2 data were from experiments in Fig. 3 at 70 °C. **(D)** Proposed model showing CdvA ring polymer has persisted following cytokinesis and is asymmetrically inherited to one of the newly divided cells.

To determine whether this behaviour of Matcha-CdvA was an artefact caused by the overexpression, we compared these data to images of formaldehyde-fixed wildtype cells stained for CdvA. The immunostained images resembled snapshots of the dynamics observed by Matcha-CdvA live imaging in that CdvA ring structures were found in constricting cells (Fig. S9A). We also observed a similar process of asymmetric CdvA partitioning using flow cytometry (Fig. S9B-E), as two distinct populations of G1 cells were observed that inherited either high or low CdvA concentrations in wildtype *S. acidocaldarius* cells (Fig. S9D) – just as had been suggested by the live cell imaging carried out using Matcha-CdvA.

Collectively, these data indicate that, while the CdvB protein ring is disassembled and degraded as cells divide, the CdvA ring is not. CdvA does not constrict to a point like CdvB1 and CdvB2, instead it constricts until it forms a small ring that lies just off-centre from the division bridge (Fig. 4A, 4D). Once the CdvA ring has constricted, the ring is then asymmetrically inherited by one of the two nascent G1 cells (Fig. 4A, 4D); similar to the way the midbody is inherited by one cell following ESCRT-III-mediated abscission in human cells.

### Practical considerations of using Matcha in live imaging

While fluorescent live cell imaging has revolutionised cell biology, significant challenges have precluded using this type of approach to image small hyperthermophiles, like *Sulfolobus*. This is important to advance our understanding of cellular processes in these organisms at high temperatures and reveal dynamic behaviours that cannot be captured by fixation-based methods.

To make live cell imaging in hyperthermophiles possible, several challenges needed to be overcome. First, a high temperature live imaging setup was required, which we and others achieved through the design of a chamber that included a heated lens collar and a heated lid (*22–24*) and selection of compatible small molecule dyes (*25, 26*). Next, since bacteria and archaea tend to be small, a high resolution imaging setup is typically desired, for which we and others used super-resolution microscopy methods including SoRa spinning disk confocal (*25*) and structured illumination microscopy (*26*). Third, while efforts have been made to make tools for live imaging of fluorescently labelled proteins, the success of this approach has been limited by low efficiency of fluorescent protein folding, decrease in fluorescent protein brightness and fast photobleaching at high temperature. These mean that, even though it is possible to label a specific protein or structure in live cells using TGP (*8*) or dye-binders (*27, 28*) it has not yet been possible to image protein dynamics during division in an extremophile like *Sulfolobus*.

Through the evolution of Matcha, a fluorescent variant of TGP that is ∼50 times brighter, we were able to image target proteins live during cell division at physiological temperature of *Sulfolobus*. A simple way to improve the fluorescent signal in these experiments is to decrease the imaging temperature to 65-70 °C (Fig. 2B, Fig. S10). As we show in this paper, *Sulfolobus* cells can still proliferate at these temperatures, albeit with a decrease in growth rate (Fig. S3). Thus, the optimal imaging temperature requires empirical testing to strike a balance between the overall growth speed and fluorescent properties such as signal-to-noise ratio and photostability. The lower fluorescent signal at high temperature also highlights the importance of optimising the photon budget when choosing the imaging modality. While confocal techniques provide improved spatial resolution and optical sectioning power compared to traditional epifluorescence microscopy, a drop in signal intensity is expected when using spinning disk confocal microscopy (*29, 30*). Wide-field imaging methods may therefore be more suitable for samples with low abundance of fluorophores to enhance sensitivity. Lastly, in our experiments, the fluorescent signal of Matcha was found to be lower when fused to the C-terminus of a target protein (Fig. S11), similar to the case of TGP (*8*). This likely indicates the lower co-translational folding efficiency of Matcha when employed as a C-terminal fluorescent tag, and N-terminal tagging is thus generally preferable for maximising fluorescence signal when functional interference is not anticipated.

### Using Matcha to reveal new cell biology

Using this approach, we have been able to reveal the dynamics of ESCRT-III ring formation, ring constriction and ESCRT-III polymer disassembly (once the polymer has constricted to a point) in living cells. Our analysis complements previous work carried out using fixed cell imaging (*15, 16, 31–33*) and live cell imaging using fluorescent membrane dyes (*16, 22*).

Much more surprisingly, however, through this analysis we have also seen that CdvA forms a template ring that is maintained throughout cytokinesis, which is shifted off to one side as the CdvB1/B2 ring constricts and is then asymmetrically inherited by one of the two daughter cells. This is reminiscent of the process seen in human cells, where the template of ESCRT-III-dependent cytokinesis, the midbody, is retained during the process of cytokinesis and then asymmetrically inherited (*34*). While functions have been ascribed to the midbodies in animal cells (*35*), the asymmetric inheritance of CdvA observed in *Sulfolobus* may simply be a way for cells to handle a physically stable structure that acts as a template for the division ring. The persistent presence of CdvA in the ring also raises the question of how CdvB is selectively removed first without apparent disassembly of CdvA/B1/B2 rings, and if CdvA provides mechanical aid as CdvB1 and CdvB2 constrict the membrane. Our results thus suggest a modification of the current cytokinesis model (Fig. 4D), implying that CdvA may have roles beyond serving as a template for division ring assembly in *Sulfolobus* and other Thermoproteota.

By studying the polymer dynamics of CdvB2 and CdvA using Matcha, we have unveiled previously unknown mechanistic steps of *Sulfolobus* cell division, demonstrating how this work can shed new light on archaeal cell biology. It is our hope that this work will stimulate further protein engineering studies in model hyperthermophiles such as *Sulfolobus* to expand the palette of fluorescent proteins. Note that there is also a recent report of using dye-binders as an alternative approach (*27*). We also hope that this will enable the development of diverse molecular tools with enhanced thermostability and folding efficiency, with potential applications extending beyond extremophile biology.

## Materials and Methods

### Cell culture and induction conditions

All *Sulfolobus acidocaldarius* strains were cultured in Brock medium as previously described (*36*) at 75 °C unless otherwise noted with constant shaking at 160 rpm. The growth media is supplemented with 0.1% NZ-amine, 0.2% sucrose, 0.04 g/L FeCl_3_(H_2_O)_6_ (here on referred to as BNS media) and adjusted to pH 3 by addition of 50% (v/v) sulfuric acid. For the uracil auxotrophic strains MW001 and MW007 (ΔSuaI) (*37*), 4 mg/L uracil was supplemented. For all experiments, the cultures were passaged at least once after inoculation from the frozen glycerol stock. For experiments with arabinose-induction of less than 8 hours, cells were grown to early log phase (OD_600_∼0.1 to 0.2) before adding L-arabinose to a final concentration of 0.2%. Prolonged induction was performed by addition of 0.2% arabinose in the media during passage and incubated overnight to OD_600_ of ∼0.2 at the indicated temperature. For storage of *S. acidocaldarius* strain, 5 to 10 mL of culture was grown to stationary phase and centrifuged at 4000 x g for 4 min. The cell pellet was then resuspended in BNS media containing 50% glycerol at pH 5 and stored at −70 °C as glycerol stock.

### Generation of mutant library in *S. acidocaldarius*

To generate the site-saturation mutant library of TGP fused with the *Saccharalobus solfataricus* β-galactosidase (LacS), BamHI restriction enzyme cutting site was first introduced to the linker region of the arabinose-inducible TGP-linker-LacS-HA plasmid (*8*) by Q5 site-directed mutagenesis (New England Biolabs). The resulting plasmid was then digested with NcoI and BamHI to remove wild type TGP, followed by gel extraction. Site saturation variant library of codon-optimized TGP with flanking NcoI and BamHI digestion sites was custom synthesized commercially (Twist Bioscience). The library contained single amino acid mutation from 48 selected sites with 47 sites passing quality control by next generation sequencing (see Table S1 for summary of mutation sites). The variant library was then split into 4 (each containing libraries from ∼12 mutation sites), digested with BamHI and NcoI followed by spin column purification. The digested libraries were then ligated with double digested vector with T4 DNA ligase and transformed to NEB 10-beta electrocompetent cells followed by antibiotic selection in liquid culture. Plasmids of site-saturation variant libraries were then purified from the transformed *E. coli* cells.

The plasmid libraries were then transformed by electroporation into MW007 competent cells, which is a markerless deletion strain of the *S. acidocaldarius* restriction enzyme SuaI *(saci_1976)* in the uracil auxotrophic strain MW001 constructed by the pop in/pop out method (*38*), at 2000V, 600Ω, 25 μF in 1 mm Gene Pulser Electroporation Cuvette (Biorad). The transformed cells were then recovered in recovery media (Brock media without supplementation of additional NZ-amine and FeCl_3_ solution, pH ∼5) for 1 hr at 75 °C, followed by inoculation into 10 times volume of pH 5 BNS media. The transformed cells were then grown at 75 °C with shaking for two days, centrifuged down and inoculated into 10 mL of pH 3 media until early stationary phase. The Sulfolobus cells containing site-saturation variant library of TGP fused with LacS-HA were then frozen as glycerol stocks and stored in −70 °C.

### Molecular Genetics and plasmid transformation

The TGP fusion protein constructs were constructed by replacing the LacS sequence in pSVAaraFX-TGP-linker-LacS-HA plasmid (*8*) with the sequence PCR amplified from the genomic DNA of *S. acidocaldarius* DSM639 through Gibson assembly. For TGPv2-linker-LacS-HA construct, dsDNA fragment containing TGPv2 codon-optimized sequence along with flanking sequences overlapped with arabinose-inducible promoter and the linker (GSAGSAAGSG) was custom synthesized (gBlock, IDT) and replace TGP coding sequence in the pSVAaraFX-TGP-linker-LacS-HA plasmid used for the generation of site-saturation variant library by Gibson assembly. For C-terminally tagged LacS constructs, LacS-linker-TGP-HA and LacS-linker-TGPv2-HA were synthesized as gBlocks and cloned into the double-digested (NcoI, XhoI) pSVAaraFX-HA vector via Gibson assembly. Constructs of LacS fused with TGPv3 and Matcha were then generated by Q5-site directed mutagenesis (New England Biolab) to incorporate point mutations sequentially. Matcha-Cren7-HA expression plasmids were generated by replacing LacS sequence from pSVAara-FX-Matcha-linker-LacS-HA plasmid with PCR amplified Cren7 *(saci_1307)* from the genomic DNA of DSM639. NcoI and BamHI digestion sites were introduced to the N-termius of Cren7 by PCR, and ligated to gel purified pSVAara-FX-Matcha-linker-LacS-HA plasmid digested with NcoI/BamHI.

Matcha-CdvA-HA overexpression construct was constructed by PCR amplification of the CdvA *(saci_1374)* open reading frame from DSM639 genomic DNA. Flanking BamHI and C-terminal HA tag with ApaI restriction sites were appended via PCR primers. The resulting PCR product was cloned via restriction digest into the gel purified, double digested (BamHI/ApaI) Matcha-LacS-HA construct to replace LacS coding sequence.

To generate the long linker version of LacS-Matcha-HA plasmid, an SGGGSGG linker sequence were inserted to flank the C-terminus of LacS in LacS-Matcha-HA plasmid using by Q5-site directed mutagenesis. To construct CdvB/CdvB1/CdvB2-Matcha-HA plasmids with long linker, SGGGSGG sequence was introduced to the C-terminus of each gene by primer design during PCR amplification of the coding sequences from the genomic DNA of DSM639 and PCR linearization of the pSVAara-FX-Matcha-linker-LacS-HA vector. Additional overlapping sequence was also introduced in these PCR primer sets to serve as Gibson assembly handles. The purified PCR products were then assembled to the linearised vector via Gibson assembly.

Cloned plasmids were transformed to ER1821 E. coli strain that contains pM.EsaBC41 plasmid for cytosine methylation. The methylated plasmids were then transformed to electrocompetent MW001 uracil auxotrophic strain of *S. acidocaldarius* by electroporation, followed by resuspension in 1 mL of recovery media and incubation at 75 °C for 30 min to 1 hr. The transformed cells were then plated on Brock medium plates containing 0.6% gelrite (Sigma-Aldrich), without uracil supplemented, for 5 to 7 days at 75 °C. Single colonies were picked and grown up in BNS media without uracil, and verified through PCR genotyping and Sanger sequencing. Positive clones were frozen as glycerol stock and stored in −70 °C.

### Immunostaining and flow cytometry

Ethanol fixation was performed by stepwise addition of cold ethanol to the cell culture samples to the final ethanol percentage of 33, 50, 70% with 5 min incubation intervals on ice. 1.5 mL of fixed cells were collected via centrifugation at 8000 x g for 3 min. Cells were then washed twice in 1 mL of phosphate-buffered saline containing 0.2% Tween-20 (Sigma-Aldrich) and 3% Bovine Serum Albumin (Sigma-Aldrich) (PBSTA). Cells were resuspended in primary antibodies in 150 μL of PBSTA, and incubated overnight with shaking (500 rpm) at room temperature. The stained cells were then washed three times in 1 mL of phosphate-buffered saline containing 0.2% Tween-20 (PBST) and resuspended in 150 μL of PBSTA containing secondary antibodies. After 2 hr incubation at room temperature with shaking, the cells were washed three times in 1 mL PBST and then resuspended in 500μL of PBST containing 2 μM Hoechst (Thermo Fisher Scientific, 62249) for labelling of DNA.

Formaldehyde fixation of *S. acidocaldarius* cells were performed by addition of formaldehyde (Sigma-Aldrich) to a final concentration of 4% and incubated at room temperature for 10 minutes, followed by three washes with phosphate buffer saline (PBS). The fixed cells were then permeabilized by resuspending the cells in 0.01% of sodium dodecyl sulfate solution in PBS, and incubated at room temperature for 20 minutes. The permeabilized cells were then washed for three times with PBSTA, followed by standard antibody staining procedure as described above. The immunostained cells were then centrifuged to a LabTek chamber (Thermofisher) pre-incubated with 1% polyethyleneimine (PEI) solution, and imaged by SoRa spinning disk confocal microscopy as previously described (*16*).

For live cell sample, cell culture was diluted with pre-heated BNS+0.2% arabinose that was filtered with 0.22 μm syringe filter to the final OD_600_ of ∼0.03. The diluted cell samples were then passed through a 70 μm cell strainer (BD Falcon). The samples were kept warm in a polystyrene box containing pre-heated metal beads until the flow cytometry analysis.

All flow cytometry experiments were performed on a BD Biosciences LSRFortessa. Laser wavelengths used are 355, 488, 561 and 640nm, in conjunction with the emission filters 450/50, 530/30, 585/15 and 670/15 respectively. For the thresholding of flow cytometer, Hoechst channel (emission filter 450/50 with 355 nm excitation) was used to identify immunostained ethanol fixed cells, while side scattering (SSC) channel was used to view unfixed Sulfolobus cells. All flow cytometry data was analysed in FlowJo v10 software. Events corresponding to single ethanol fixed cells were selected by gating the diagonal population in the area vs. height plot in the Hoechst channel. For live cell samples, green fluorescence positive cells from either TGP-linker-LacS-HA or Matcha-linker-LacS-HA expressing strain were first selected and backgated to forward scattering (FSC) and SSC plot. The live cell population was then identified by applying this gate (on the FSC and SSC plot) to all recorded events.

### Cell sorting and mutant identification

*S. acidocaldarius* cultures induced by 0.2% arabinose overnight were centrifuged at 5000 x g for 3 min at 60 °C, and resuspended in warm BNS media+0.2% arabinose filtered with 0.22 μm syringe filter to OD_600_∼0.03. The diluted cells were then passed through the 70 μm cell strainers twice and kept in the polystyrene box containing pre-heated metal beads before cell sorting. Cell sorting was performed on the ThermoFisher Bigfoot and BD FACSAria Fusion with a 70 μm nozzle at 60psi. Populations of cells were identified by back gating on positive green fluorescence signal on the FSC and SSC detectors, both set to a logarithmic scale. A negative and buffer only controls were run to assess true signal above background. Both sorters used a 488nm laser excitation and emission was detected using a 520/35 BP filter on the Bigfoot and a 530/30 BP filter on the Fusion. 600,000 cells with high (∼top 5-10%), medium (only in the third round of sorting), and low green fluorescence intensity were purified and collected into 3 mL of BNS media (pH 3) at 37 °C. The sorted cells were then centrifuged at 5000 x g, 3 min at room temperature and ∼3.3 mL of supernatant was carefully removed. The pelleted cells were resuspended using the residual solution and inoculated into 10 mL of BNS media (pH 3) to grow at 75 °C for 1-2 days until clear growth. The culture was then centrifuged and resuspended with fresh 25 mL BNS media and grown at 75 °C for 1-2 days until late log phase or early stationary phase before stored as frozen glycerol stocks.

Cultures were grown from these glycerol stocks following the aforementioned culturing procedure and induced with 0.2% arabinose at 75 °C overnight. The green fluorescence signal was then examined by flow cytometry analysis. The four transformed *S. acidocaldarius* libraries were sorted separately for the first two round and were pooled together with an equal cell density for the last round of sorting. After the third round of sorting, cultures originated from the top green fluorescence signal population no longer showed higher green fluorescence intensity compared to the ones sorted from the low green fluorescence signal population, suggesting that there was no further enrichment in beneficial mutants. To identify the enriched mutations, log-phase culture from the last round of cell sorting was used as template for PCR amplifications by Q5 polymerase (New England Biolab) using two pairs of primers to cover the full region containing all mutations (Amp-seq-F1: 5’-TGGCGGCAAGTGTAATAAAGC, Amp-seq-R1: 5’-CATTACAGGTCCA TTTGGAGG; Amp-seq-F2: 5’-CAAGCAAGCATTTCCAGAAGGA, Amp-seq-R2: 5’-ATCCTCCACCACCACTACCA). The PCR amplicons were purified by PCR cleanup and sent for amplicon sequencing and SNP detection (Genewiz, Azenta Life Science). The relative abundance of each mutant is shown in Table S1.

### Live imaging

Live-cell imaging at high temperatures was performed as previously described(*22, 25*) using the in-house built heating elements coupled with the objective heater (Okolab) on the Nikon Ti2 Eclipse microscope (the ‘Sulfoscope’ setup described in (*22*)). Briefly, 25 mm diameter coverslips were mounted on imaging chambers (Attofluor, Invitrogen A7816) and filled with 400 µL BNS media, pH 3. The medium was dried on the coverslip at 75 °C, and then the chambers were washed with BNS and placed in the preheated Sulfoscope to equilibrate to the desired temperature (65, 70 or 75 °C). 10 mL *S. acidocaldarius* culture at OD_600_∼0.15 was maintained at the set temperature and stained with CellMask Deep Red Plasma Membrane (Invitrogen C10046; 1:5000). 450 µL of the stained cell suspension was transferred to the preheated imaging chamber, and cells were immobilised using semi-solid Gelrite pads (BNS-Gelrite: 0.6% Gelrite, 1× Brock medium pH 5, 20 mM CaCl_2_). To prepare immobilisation pads, 15 mL of the melted BNS-Gelrite solution was added to 9 cm petri dishes and allowed to solidify at room temperature. Half-moon pieces cut with a 7-mm circle punch were placed onto 13-mm coverslips. Immediately before imaging, pads were incubated at 75 °C for ∼5 min to dry, allowing the edges to curve. Cells were imaged at the concave edge, where mobility is restricted while limiting mechanical compression from the pad. Images were acquired on a Nikon Eclipse Ti2 inverted microscope equipped with a Yokogawa SoRa scanner unit with an additional 2.8x and 4x magnifications and Prime 95B scientific complementary metal-oxide semiconductor (sCMOS) camera (Photometrics). Imaging was performed with a 60x oil immersion objective (Plan Apo 60x/NA 1.45, Nikon) using a custom formulated immersion oil for high temperature imaging (maximum refractive index matching at 70 °C, n =1.515 ± 0.0005; Cargille Laboratories). Time-lapse imaging was performed at 1 frame per min (75, 70 °C) or 3 min interval (65 °C) with at least 3 biological replicates for each strain. Image analyses were performed using Fiji (*39*).

### Quantitative fluorescence image analysis

Unprocessed images were analysed using ImageJ (2.16.0). To quantify the CdvB2 cytosolic fluorescence fraction, specific regions of interests (ROI) were measured and averaged at three different division stages (pre-ring, pre-constriction and post-abscission). The regions were chosen to avoid any polymeric fraction. Three frames were averaged for each ROI and the background, measured from a region outside the cell, subtracted. For each cell, the values were normalized to the pre-constriction (i.e., full ring assembly) stage.

Time-resolved measurements of cytosolic CdvB2 were performed using an ROI avoiding CdvB2 polymers in all analysed frames. Twenty frames before and after the end of observable cell constriction (t=0) were quantified. The signal was then background subtracted and normalized to average of intensity at t= −1 to +1 min (3 frames).

The symmetry score of CdvA and CdvB2 was determined by measuring average matcha fluorescence intensity in the two daughter cells. The signal was then background subtracted. Finally, the symmetry score was defined as the ratio of the lower to higher signal intensity between the two daughter cells.

### Assessment of growth temperature effect on *S. acidocaldarius*

Wild type (strain DSM639) *S. acidocaldarius* cells were used to assess the growth speed and percentage of cells in the division phase at different temperatures. Starting culture of wild type cells were first grown at 75 °C to OD_600_ ∼0.4. The cells were then inoculated to pre-heated cultures at 75, 70 or 65 °C respectively to grow for 19-20 hours at the corresponding temperatures with shaking. The final OD_600_ was recorded. The apparent doubling time was calculated as *Growth time*/ log_2_*(OD*_*final*_ /*OD*_*initial*_).

To estimate the percentage of dividing cells, DSM639 cells grown at different temperatures were ethanol fixed at OD_600_ ∼ 0.2, followed by immunostaining against CdvB and CdvB1. The immunostained cells were then analysed by flow cytometry as previously described (*40*). Cells in the pre-constriction phase (high CdvB, high CdvB1 signals with 2N DNA content) and constricting phase (CdvB negative, high CdvB1 signal with 2N DNA content) were then quantified using FlowJo (v10.10) as previously described (*9*).

### Expression and purification of recombinant proteins

To generate TGP and Matcha (both with C-terminal His_6_-tag) expression plasmids, pET22b plasmid was digested with restriction enzymes NdeI and XhoI, followed by gel extraction. *E. coli* codon optimized TGP and Matcha with flanking overlapping sequences with the doubled digested pET22b vector were custom synthesized (Integrated DNA Technologies gBlocks), and cloned into the digested pET22b vector by Gibson assembly. To express recombinant TGP and Matcha, pET22b-TGP-His_6_ and pET22b-Matcha-His_6_ were transformed to *E. coli* BL21-DE3(pLysS) (Invitrogen), and grown to OD_600_ ∼0.4-0.6, followed by induction of 0.5 mM IPTG at 37 °C for 3.5 hr with constant shaking. The *E. coli* cells were harvested, frozen and stored in −70 °C until purification.

To purify the recombinant TGP or Matcha, the cell pellet was thawed on ice and resuspended in lysis buffer containing 30 mM NaH_2_PO4, 150 mM NaCl, 10 mM imidazole pH 6.5 supplemented with 0.5 mM DTT, 0.1% TritonX-100, cOmplete protease inhibitor cocktail (EDTA free) and 0.06 U/μL Benzonase (Sigma). The cells were sonicated on ice for cell lysis. The crude lysate was clarified by centrifugation and incubated with Ni-NTA agarose resin (Qiagen) in the cold room with agitation for at least 20 min. The lysate and resin were then poured into a gravity column, and the resin bound with recombinant fluorescent proteins were then washed with at least 10 column volume of lysis buffer, followed by stepwise elution of increase imidazole concentration (50, 100, 200, 300, 500 mM) at room temperature. Peak fractions were heated at 65 °C for 15 minutes to induce aggregation of contaminating proteins from E. coli, followed by centrifugation at 20,000 x g, 15 min at room temperature. The supernatant was pooled and concentrated to ∼500 μL with a centrifugal filter and further purified with a size exclusion column, eluted with phosphate buffer (30 mM NaH_2_PO4, 150 mM NaCl, 1 mM DTT pH 6.5). The purified fractions were pooled and stored at either 4 °C or flash frozen with liquid nitrogen and stored as small aliquots in −70 °C. Concentration of the purified proteins was determined by Bradford assay (Pierce) using bovine serum albumin (BSA) as standard.

### *In vitro* fluorescent assays and spectral measurements

Absorption spectra were obtained using Cary 50 Bio UV-Vis spectrophotometer (Varian) with 16 μM of the target fluorescent protein in phosphate buffer pH 6.5 at room temperature. Fluorescence emission spectra were measured with Cary Eclipse fluorimeter (Varian) using 4 μM of the target fluorescent protein with excitation at 485 nm.

Thermal melting assay was performed using the ViiA7 Real-Time PCR system (Thermofisher). 0.5 μM of green fluorescent proteins (EGFP, TGP, Matcha) in phosphate buffer (pH 6.5) supplemented with 1 mM DTT was first incubated at 25 °C for 3 min, followed by a constant heat ramp (0.015 °C/s) to 95 °C. The fluorescence signal was normalized by the intensity at 25 °C recorded at the start of the heating ramp. Isothermal melting assay was performed similarly but the samples were heat up quickly (1.6 °C/s) and hold at 83 °C with the fluorescence signal recorded with 1 min interval. The fluorescence signal was then normalized by the first time point (T=1 min) after heating to 83 °C.

For chemical treatment assay, various concentrations of buffered guanidine HCl solution (in phosphate buffer pH 6.5) was mixed with 300 nM of each green fluorescent protein tested in a 96 well plate. The plate was sealed and incubated in the dark at room temperature overnight to reach equilibrium denaturation. The fluorescence signal was then measured using the PHERAstar FSX microplate reader (BMG Labtech) with 485 nm excitation and 520 nm emission. Hydrogen peroxide solution was prepared similarly in phosphate buffer pH 6.5 and quickly mixed with each fluorescent proteins using a multichannel pipette. The plate was incubated at room temperature for 20 minutes and the fluorescence signal recorded by the microplate reader using the same setting. Fluorescence intensity of untreated TGP and Matcha (1 μM each) was measured under the same condition using the microplate reader.

## Acknowledgement

We would like to thank Dr. Phil Holliger, Dr. Niklas Freund, Dr. Edoardo Gianni and Carlos Piedrafita Alvira for the advice on strategies of directed evolution, and Dr. Sherman Foo and Dr. Ganesh Agam for the discussion. We would also like to thank Dr. James Manton for the assistance on data analysis and discussion. All flow cytometry and cell sorting experiments were performed at the Medical Research Council Laboratory of Molecular Biology Flow Cytometry core facility, and we would like to thank members of the Flow Cytometry Facility for their technical support. We would also like to thank Dr. Stephen McLaughlin and the Biophysics Facility of MRC-Laboratory of Molecular Biology for the assistance with biophysical characterizations. Y.-W.K. was supported by an EMBO postdoctoral fellowship (ALTF 903-2021) and by the Medical Research Council-Laboratory of Molecular Biology (MC_UP_1201/27); A.R.-M. is supported by an EMBO postdoctoral fellowship (ALTF 431-2025); T.D. is supported by the Royal Society (URF\R1\221086, RF\ERE\221078). A.C. was funded by an EMBO Postdoctoral Fellowship (ALTF 1041-2021) and a Marie Skłodowska-Curie Individual Fellowship (101068523) provided by UKRI; B.B. received support for work in Sulfolobus from the Medical Research Council-Laboratory of Molecular Biology (MC_UP_1201/27), the Wellcome Trust (222460/Z/21/Z), and the Life Sciences Moore-Simons Foundation (735929LPI).

## Competing interests

The authors declare that they have no conflict of interest.

## Author contributions

Y.-W.K. and B.B. conceived the project; Y.-W.K. performed all engineering and characterization of Matcha; Y.-W.K., A.R.-M., T.D. generated and characterized cell strains; A.R.-M. and T.D. performed imaging experiments with preliminary data collected by Y.-W.K. and A.C.; F.Z. and P.A.P. performed the cell sorting experiment and provided technical support for flow cytometry; M.W. and S.A. generated the cell line used in directed evolution; G.D. made initial observation of asymmetric inheritance of CdvA from fixed cells; B.B. supervised the project; Y.-W.K. and B.B. wrote the manuscript with input from all authors.

## Notes

### Competing Interest Statement

The authors have declared no competing interest.

