## Supplemental materials for "A green fluorescent protein for live imaging in hyperthermophiles"

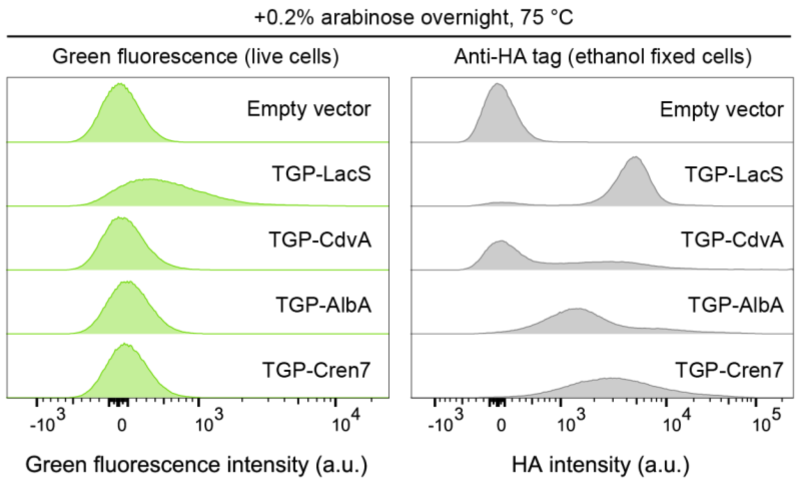


**Figure S1. TGP fusion proteins barely fluoresce above background levels at 75 °C after overnight induction in *S. acidocaldarius*.** Example flow cytometry histograms of live (left) and immunostained fixed cells (right) expressing LacS, CdvA, AlbA, Cren7 fusion proteins (N-terminal TGP and C-terminal HA tag) respectively showed that only strain expressing LacS fusion displayed a population with positive green fluorescence above the background (left), while all other fusion proteins were expressed (right) without discernible green fluorescence (left).

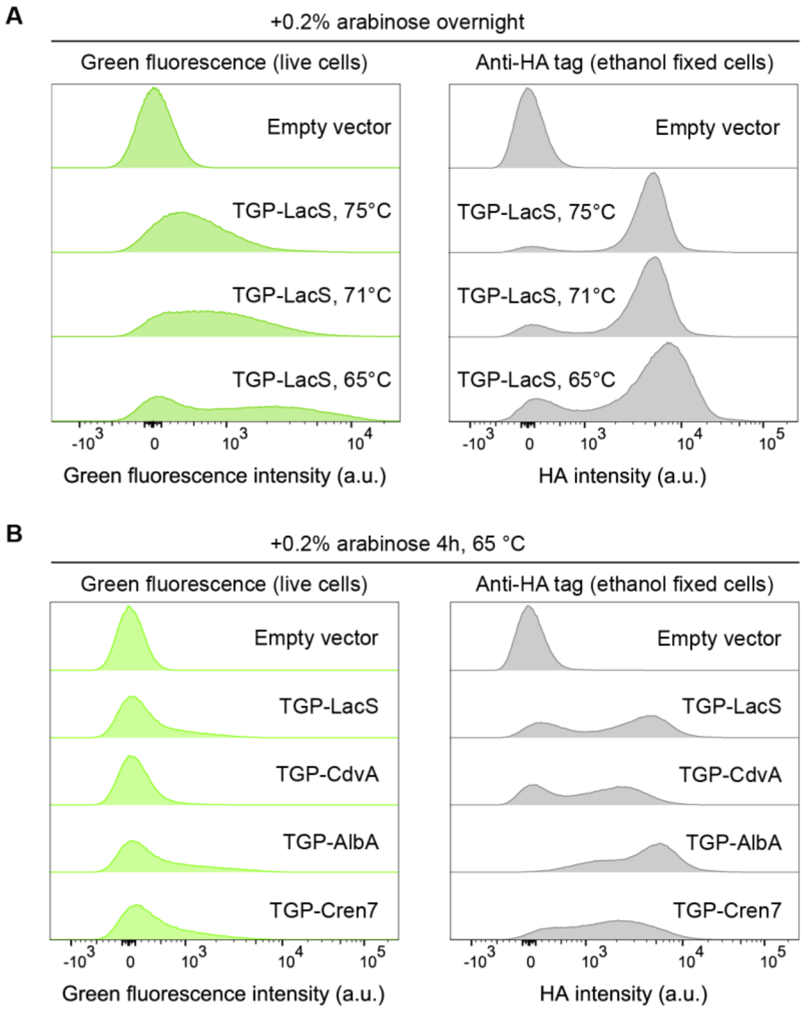


**Figure S2. TGP-fusion protein fluorescence was visible at levels marginally above background at temperatures below 75°C. (A)** Example flow cytometry histograms of cells expressing TGP-LacS fusion protein overnight from 75 to 65 °C showed increasing green fluorescence signal in the live cells (left) at lower temperature, with similar expression level (HA signal of immunostained fixed cells, right). **(B)** All TGP fusion proteins showed limited green fluorescence signal (live cell, left) at 65 °C after 4 hr of arabinose induction despite clear induction (HA signal of immunostained fixed cells, right).

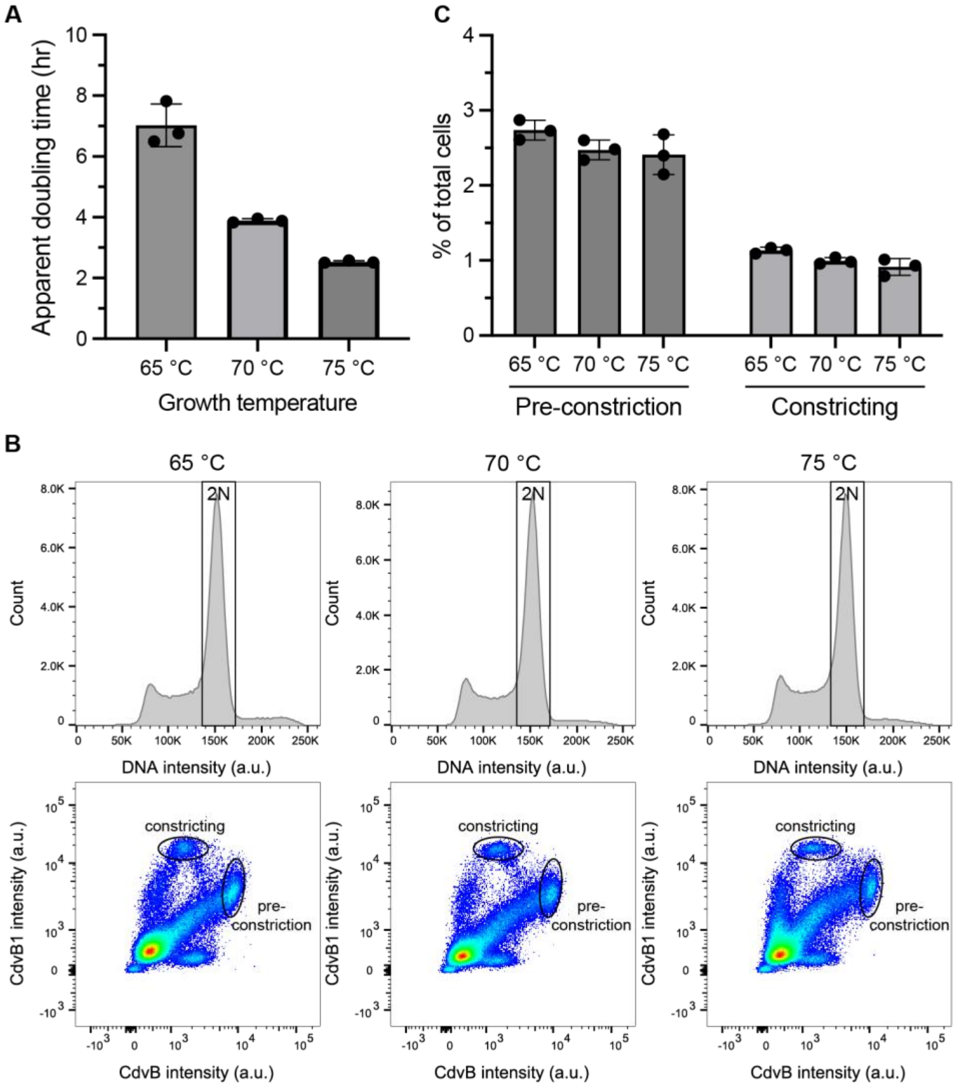


**Figure S3. Effects of growth temperature on wildtype *S. acidocaldarius* (DSM639). (A)** Decrease in growth temperature increases the apparent doubling time. **(B)** Example flow cytometry DNA histograms (top) and 2D plots of CdvB vs CdvB1 staining (bottom) of immunostained asynchronous DSM639 cultures (ethanol fixed) grown at different temperatures. **(C)** Percentages of cells in pre-constriction and constricting phase are similar in asynchronous DSM639 cultures grown at different temperatures.

| AA Position | WT AA | Most common mutant | Sum mutation frequency | A | C | D | E | F | G | H | I | K | L | M | N | P | Q | R | S | T | V | W | Y | DEL |
|---|---|---|---|---|---|---|---|---|---|---|---|---|---|---|---|---|---|---|---|---|---|---|---|---|
| 12 | I | n/a | | | | | | | | | WT | | | | | | | | | | | | | |
| 16 | M | n/a | | | | | | | | | | | | WT | | | | | | | | | | |
| 24 | K | E | | | | | | | | | | WT | | | | | | | | | | | | |
| 28 | E | K | | | | | WT | | | | | | | | | | | | | | | | | |
| 32 | I | Y | | | | | | | | | WT | | | | | | | | | | | | | |
| 34 | K | W | | | | | | | | | | WT | | | | | | | | | | | | |
| 37 | E | D | | | | | WT | | | | | | | | | | | | | | | | | |
| 42 | L | I | | | | | | | | | | | WT | | | | | | | | | | | |
| 55 | S | n/a | | | | | | | | | | | | | | | | | WT | | | | | |
| 56 | Y | F | | | | | | | | | | | | | | | | | | | | | WT | |
| 59 | L | n/a | | | | | | | | | | | WT | | | | | | | | | | | |
| 61 | P | n/a | | | | | | | | | | | | | | WT | | | | | | | | |
| 62 | A | n/a | | WT | | | | | | | | | | | | | | | | | | | | |
| 63 | F | L | | | | | | WT | | | | | | | | | | | | | | | | |
| 68 | R | n/a | | | | | | | | | | | | | | | | WT | | | | | | |
| 69 | A | V | | WT | | | | | | | | | | | | | | | | | | | | |
| 71 | T | C | | | | | | | | | | | | | | | | | | WT | | | | |
| 75 | E | P | | | | | WT | | | | | | | | | | | | | | | | | |
| 76 | D | n/a | | | | WT | | | | | | | | | | | | | | | | | | |
| 78 | P | V | | | | | | | | | | | | | | WT | | | | | | | | |
| 84 | A | n/a | | WT | | | | | | | | | | | | | | | | | | | | |
| 96 | T | I | | | | | | | | | | | | | | | | | | WT | | | | |
| 97 | Y | F | | | | | | | | | | | | | | | | | | | | | WT | |
| 102 | I | K | | | | | | | | | WT | | | | | | | | | | | | | |
| 105 | A | V | | WT | | | | | | | | | | | | | | | | | | | | |
| 107 | S | n/a | | | | | | | | | | | | | | | | | WT | | | | | |
| 111 | M | L | | | | | | | | | | | | WT | | | | | | | | | | |
| 117 | F | I | | | | | | WT | | | | | | | | | | | | | | | | |
| 118 | Y | C | | | | | | | | | | | | | | | | | | | | | WT | |
| 120 | I | n/a | | | | | | | | | WT | | | | | | | | | | | | | |
| 121 | R | H | | | | | | | | | | | | | | | | WT | | | | | | |
| 141 | W | n/a | | | | | | | | | | | | | | | | | | | | WT | | |
| 144 | S | n/a | | | | | | | | | | | | | | | | | WT | | | | | |
| 145 | T | R | | | | | | | | | | | | | | | | | | WT | | | | |
| 149 | Y | n/a | | | | | | | | | | | | | | | | | | | | | WT | |
| 157 | G | n/a | | | | | | | WT | | | | | | | | | | | | | | | |
| 159 | V | n/a | | | | | | | | | | | | | | | | | | | WT | | | |
| 161 | M | n/a | | | | | | | | | | | | WT | | | | | | | | | | |
| 165 | L | n/a | | | | | | | | | | | WT | | | | | | | | | | | |
| 169 | G | R | | | | | | | WT | | | | | | | | | | | | | | | |
| 177 | T | n/a | | | | | | | | | | | | | | | | | | WT | | | | |
| 192 | E | F | | | | | WT | | | | | | | | | | | | | | | | | |
| 195 | H | n/a | | | | | | | | WT | | | | | | | | | | | | | | |
| 198 | E | V | | | | | WT | | | | | | | | | | | | | | | | | |
| 211 | L | n/a | | | | | | | | | | | WT | | | | | | | | | | | |
| 212 | Y | R | | | | | | | | | | | | | | | | | | | | | WT | |
| 214 | H | n/a | | | | | | | | WT | | | | | | | | | | | | | | |

High

Low

**Table S1. Relative abundance of each mutation site after directed evolution.** Mutation sites included in Matcha is highlighted in blue.

```
Matcha   1   MAASVIKPEMKIKLRMEGAVNGHKFVIKGEGIGKPYEGTQTIDLTVEEGA   50
             MAASVIKPEMKIKLRMEGAVNGHKFVI+GEGIGKPYEGTQT+DLTVEEGA
   TGP   1   MAASVIKPEMKIKLRMEGAVNGHKFVIEGEGIGKPYEGTQTLDLTVEEGA   50

Matcha  51   PLPFSFDILTPAFQYGNRAFTKYPEDIPDYFKQAFPEGYSWERSMTYEDQ  100
             PLPFS+DILTPAFQYGNRAFTKYPEDIPDYFKQAFPEGYSWERSMTYEDQ
   TGP  51   PLPFSYDILTPAFQYGNRAFTKYPEDIPDYFKQAFPEGYSWERSMTYEDQ  100

Matcha 101   GKCIATSDITMEGDCFIYEIRFDGTNFPPNGPVMQKKTLKWEPSTEKMYV  150
             G CIATSDITMEGDCF YEIRFDGTNFPPNGPVMQKKTLKWEPSTEKMYV
   TGP 101   GICIATSDITMEGDCFFYEIRFDGTNFPPNGPVMQKKTLKWEPSTEKMYV  150

Matcha 151   EDGVLKGDVEMALLLEGGGHYRCDFKTTYKAKKDVRLPDAHFVDHRIVIL  200
             EDGVLKGDVEMALLLEGGGHYRCDFKTTYKAKKDVRLPDAH VDHRI IL
   TGP 151   EDGVLKGDVEMALLLEGGGHYRCDFKTTYKAKKDVRLPDAHEVDHRIEIL  200

Matcha 201   SHDKDYNKVRLYEHAEARYSGGGSGGG  227
             SHDKDYNKVRLYEHAEARYSGGGSGGG
   TGP 201   SHDKDYNKVRLYEHAEARYSGGGSGGG  227
```

**Figure S4. Alignment of amino acid sequences of Matcha and TGP.** Mutations introduced were highlighted in blue boxes.

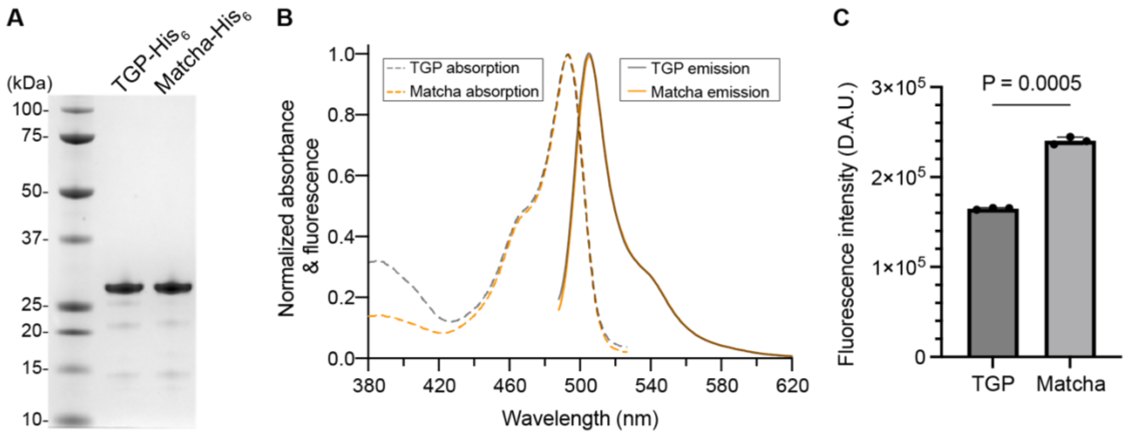


**Figure S5. Although Matcha is 50X brighter *in vivo*, TGP and Matcha have similar spectral properties *in vitro*. (A)** SDS-PAGE gel of purified TGP and Matcha with C-terminal $His_6$-tag expressed in *E. coli* recombinantly. **(B)** Absorption and emission spectra of TGP and Matcha at room temperature. **(C)** Average fluorescence intensity of 1 μM TGP- $His_6$ and 1 μM Matcha- $His_6$ measured at room temperature. Welch t-test, N=3 replicates. Consistent intensity increase of Matcha compared to TGP was observed in two different purification batches.

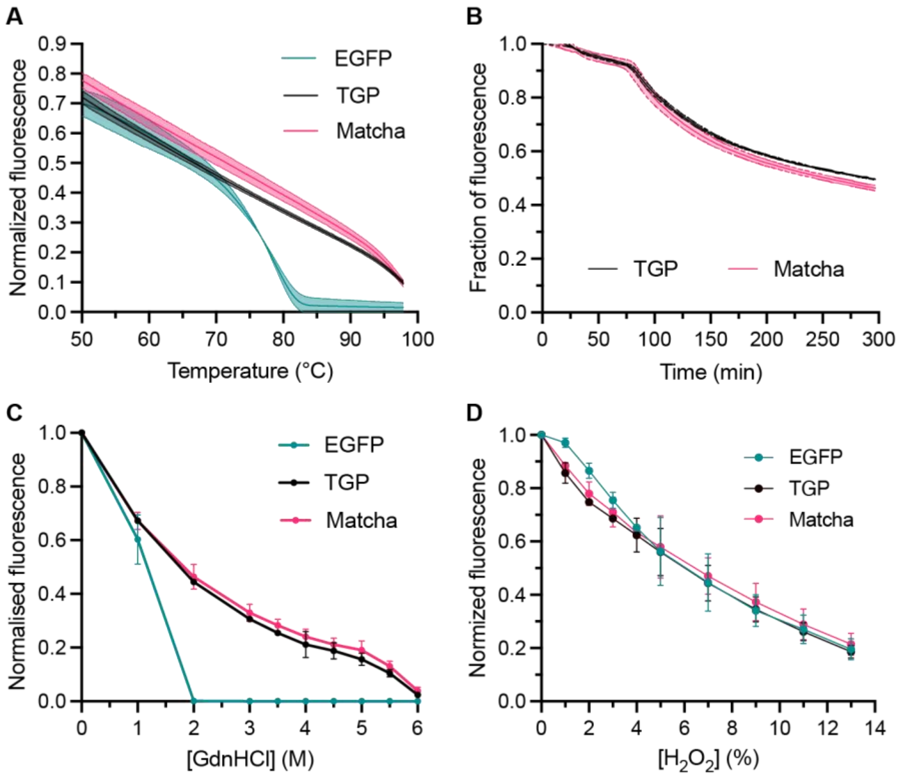


**Figure S6. TGP and Matcha have similar thermal and chemical stability profiles *in vitro*. (A)** Thermal melting assay of purified EGFP, TGP and Matcha. Mean ± 95% CI. N=3 replicates. Fluorescence signal was normalized by intensity at 25 °C immediately prior to the heating ramp. **(B)** Isothermal melting of TGP and Matcha at 83 °C with signal normalized to the first time point (time=1 min). Mean ± SD. N=3 replicates. **(C)** Equilibrium unfolding of EGFP, TGP and Matcha with buffered guanidine hydrochloride (GdnHCl) at room temperature. Fluorescence intensity was normalized to the buffer control (0 M GdnHCl). Mean ± SDs; N=3 replicates. **(D)** Fluorescence intensity of EGFP, TGP and Matcha treated by various concentrations of hydrogen peroxide for 20 min at room temperature. Fluorescence intensity was normalized by the buffer control (0% $H_2O_2$). Mean ± SDs; N=3 replicates.

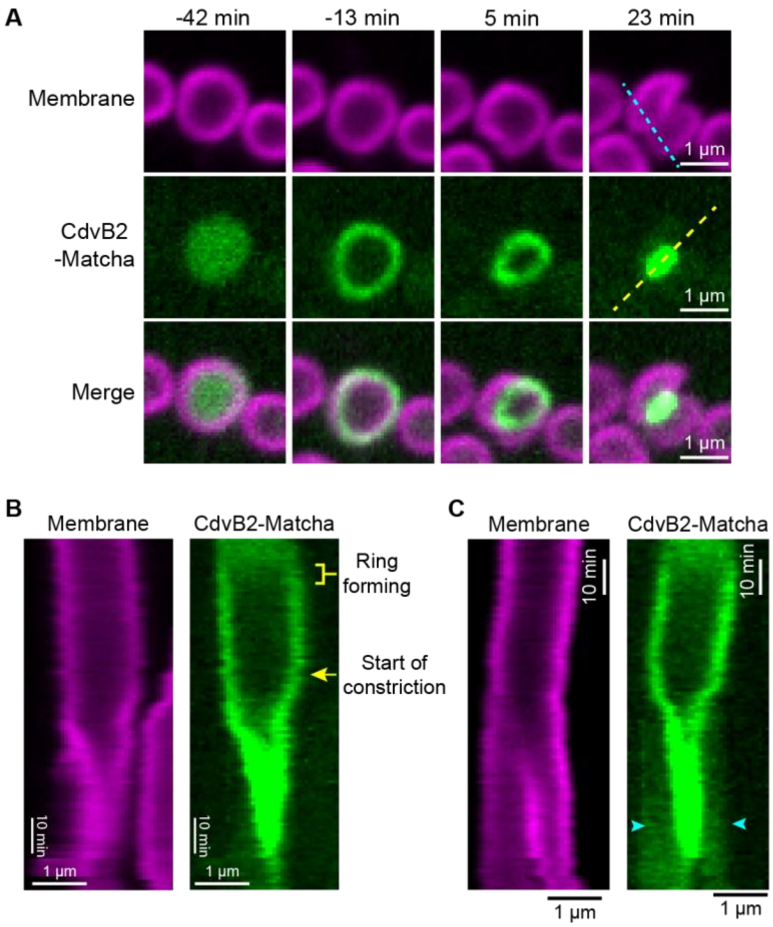


**Figure S7. Example of CdvB2-Matcha ring assembly and disassembly. (A)** Example timelapse images of top-view Matcha-CdvB2 ring formation and constriction at 70 °C (4 hr arabinose induction). The initiation of membrane constriction was set to time = 0 min. **(B)** Kymograph along the dashed yellow line in (A), showing the CdvB2-Matcha ring formation (bracket) and start of ring constriction (arrow). **(C)** Kymograph along the cyan dashed line. The higher cytosolic CdvB2-Matcha signal was visible close to the later phase of ring constriction (arrowheads), indicating the disassembly of CdvB2 ring polymer.

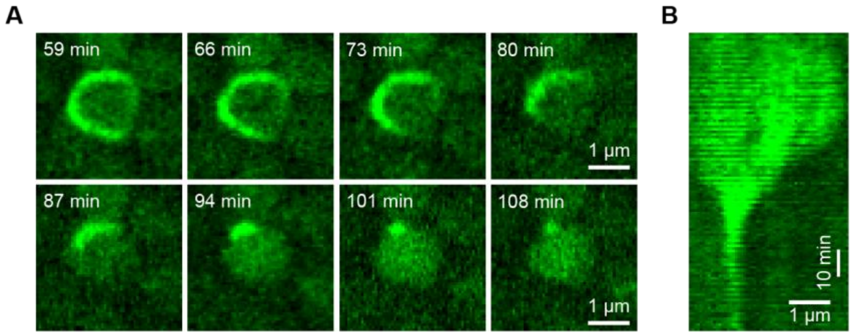


**Figure S8. Example of CdvB2-Matcha polymer shrinkage in live cell imaging. (A)** Example timelapse montage showing partial CdvB2-Matcha ring depolymerization from both ends at 70 °C. The depolymerization of partial CdvB2 ring may be resulted from a failed division caused by the mechanical compression from the gel pad used for live imaging. **(B)** Corresponding kymograph of CdvB2-Matcha polymer depolymerization in (A) with line along the cell periphery.

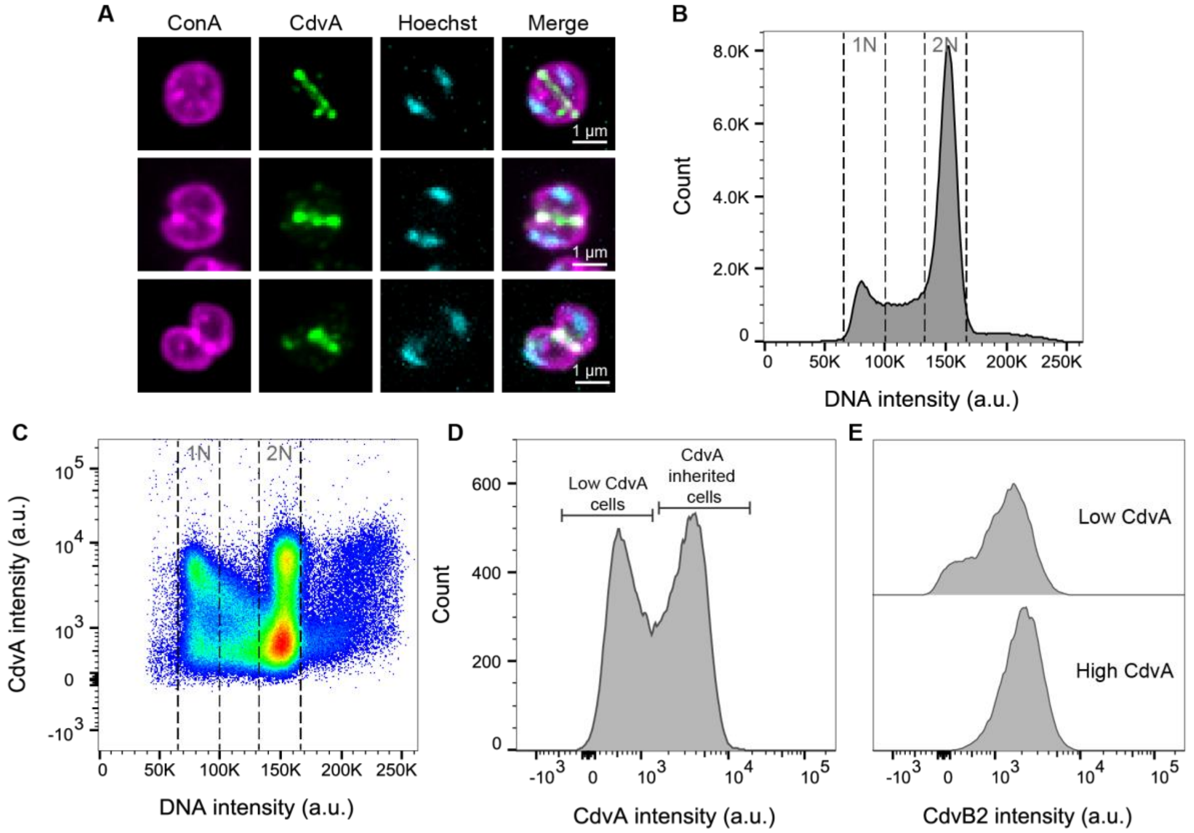


**Figure S9. Wildtype (DSM639) *S. acidocaldarius* cells show CdvA ring during cytokinesis and asymmetric inheritance of CdvA after cell division. (A)** Example immunofluorescence images (maximum projection) of formaldehyde-fixed wildtype cells synchronised by acetic acid (100 min post release), showing CdvA ring in both pre-constriction (top) and constricting phase (middle and lower panels). Note that we found CdvA rings were mostly not preserved in ethanol fixed cells, and formaldehyde fixation is needed to reduce the alteration of CdvA ring structures. Cell surface and DNA were visualised by Concanavalin A (ConA) and Hoechst staining, respectively. **(B)(C)** Example flow cytometry histogram (B) and scatter plot (C) of ethanol-fixed asynchronized wildtype cells. **(D)** CdvA intensity histogram of cells with 1N DNA content in (B) showing there are two populations of G1 cells with high and low CdvA intensity individually, indicating asymmetric inheritance after completion of cytokinesis. **(E)** CdvB2 intensity histograms of the populations of G1 cells with low and high CdvA intensity, indicating that CdvB2 inheritance has no pronounced asymmetry.

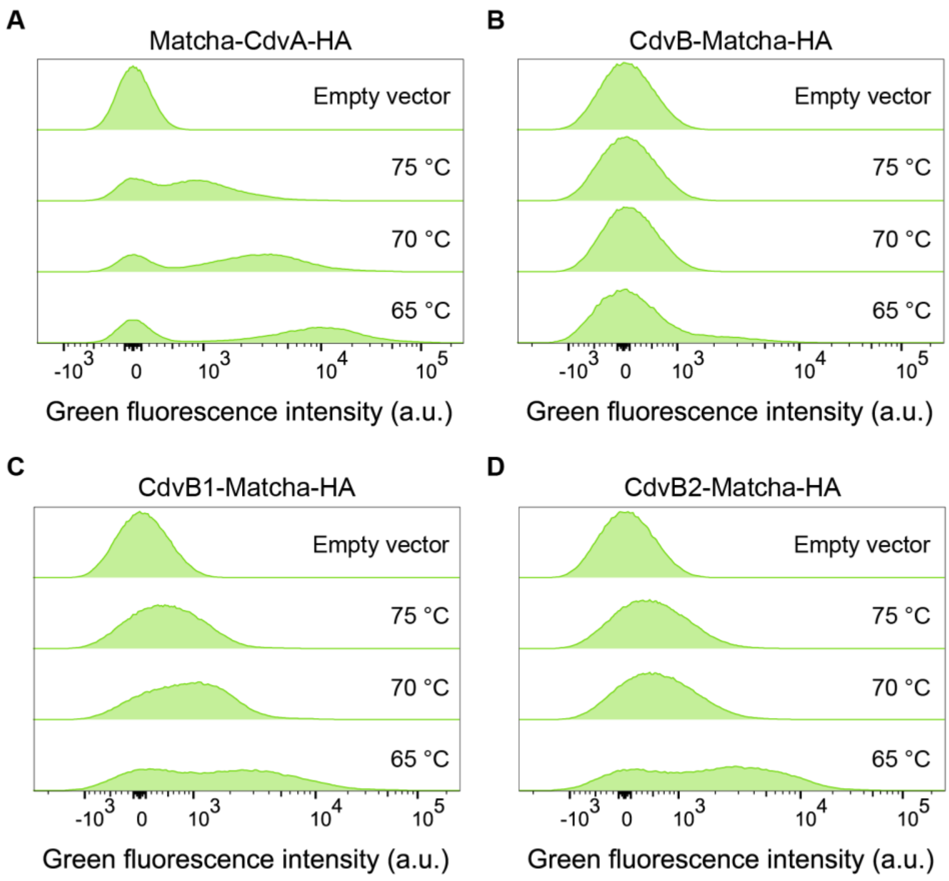


**Figure S10. Fluorescence intensity of cells expressing Matcha-tagged proteins at different temperatures. (A-D)** Example flow cytometry histograms of live cells expressing Matcha-CdvA (A), CdvB-Matcha (B), CdvB1-Matcha (C) and CdvB2-Matcha (D) induced with 0.2% arabinose for 4 hr.

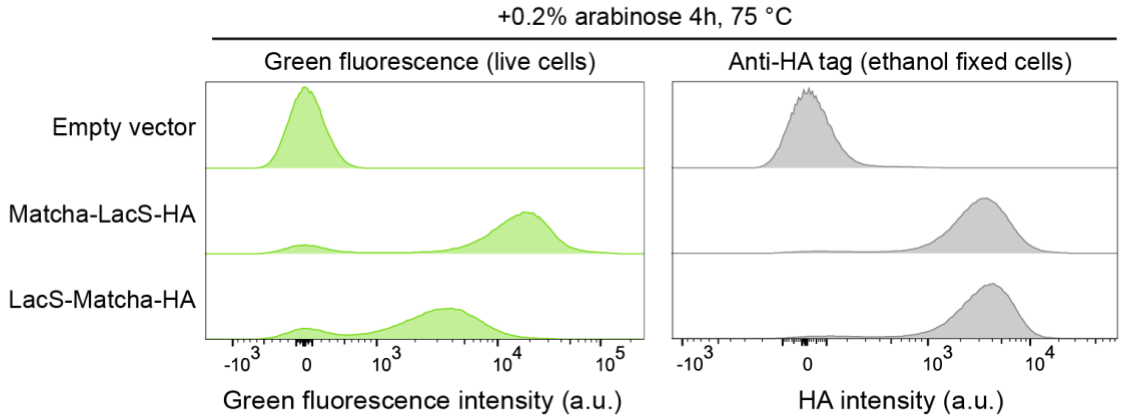


**Figure S11. Matcha shows higher fluorescent signal as an N-terminal fusion tag.** Example flow cytometry histograms of green fluorescence signal from live cells (left) and anti-HA tag signal from immunostained fixed cells (right). Note that cells expressing Matcha-LacS-HA (i.e., Matcha as an N-terminal fusion tag) showed stronger green fluorescence than cells expressing LacS-Matcha-HA, while with similar expression level based on HA tag staining signals. Note that in the LacS-Matcha-HA construct, an SGGGSGG sequence was appended to the C-terminus of LacS so that same length of flexible linker was used for both N- and C-terminal tagging constructs.
